# Stable isotope tracing of sex steroids reveals tissue-specific steroid pharmacokinetics and metabolism in intact mice

**DOI:** 10.64898/2025.12.16.694587

**Authors:** Frank Giton, Audrey Der Vartanian, Noalig Wyckens, Ioana Ferecatu, Mélanie Chester, Céline Héraud, Amine Isik, Chantal Mathis, Céline J Guigon

## Abstract

**Background and purpose:** Sex steroids play central roles in endocrine regulation and are major therapeutic targets in multiple hormone-dependent disorders. However, understanding the tissue-specific pharmacokinetics, distribution and metabolism of these hormones remains challenging *in vivo*, as most experimental approaches rely on surgical castration, which profoundly disrupts endocrine homeostasis and limits translational relevance for pharmacological studies. We therefore developed a strategy enabling quantitative analysis of steroid dynamics in physiologically intact animals.

**Experimental approach:** We established a mass spectrometry-based platform combining systemic administration of deuterium-labelled sex steroids (E2-d_4_, Testo-d_3_ and DHEA-d_5_) with high-sensitivity GC-MS/MS to simultaneously quantify exogenous tracers, endogenous steroids and their metabolites in serum and multiple tissues of intact, non-castrated mice.

**Key results:** This approach enabled temporally resolved tracking of steroid uptake, distribution and biotransformation in gonadally intact animals, providing a baseline that is closer to native physiology than in castration-based paradigms. In male and female mice, the method revealed organ-specific accumulation patterns of the metabolites of injected deuterium-labelled sex steroids, including E2-to-E1 conversion in ovary and hippocampus and Testo-to-DHT conversion in prostate and seminal vesicle, that are consistent with the distribution of steroidogenic enzymes in these tissues.

**Conclusion and implications:** This methodology provides a physiologically relevant framework for investigating steroid pharmacokinetics and pharmacodynamics *in vivo*, in which local versus systemic contributions can be dissected in future perturbation studies. By enabling quantitative mapping of steroid metabolism across tissues, it offers a powerful tool for endocrine pharmacology and for the development and evaluation of therapies targeting steroid pathways. It may, thus, facilitate mechanistic studies and preclinical evaluation of anti-androgen therapies, steroidogenesis inhibitors, and other treatments used in hormone-dependent cancers.

## Introduction

Sex steroids such as estradiol (E2), testosterone (Testo), and dehydroepiandrosterone (DHEA) play pivotal roles in regulating a wide array of physiological processes, including reproduction, metabolism, and brain function, with great variations in their levels with sex and age ^1,2^. Precise knowledge of their tissue-specific distribution, metabolic conversion, and dynamic regulation is essential for understanding both normal physiology and pathologies, including hormone-related diseases, neurodegenerative disorders and cardio-vascular diseases ^3–5^. However, elucidating these parameters *in vivo* remains a significant challenge due to the complexity of steroid metabolism and the interference from endogenous hormone pools.

Traditional experimental approaches often rely on surgical castration or gonadectomy to suppress endogenous sex steroids, enabling the study of exogenous hormone dynamics without background interference ^6,7^. While effective, these methods drastically alter the endocrine environment, leading to widespread physiological and behavioural disruptions. Numerous studies have shown that castration induces profound changes in metabolic homeostasis, immune function, neuroendocrine signalling and brain circuitry, as well as alterations in stress responses, social and sexual behaviours, and overall energy balance ^8–10^. These systemic modifications not only obscure the native roles of sex steroids but also introduce secondary effects that complicate the interpretation of downstream biological processes. Consequently, there is a critical need for approaches that allow precise tracking of steroid dynamics without imposing such extensive physiological perturbations.

To the best of our knowledge, only a few methodological frameworks currently enable quantitative distinction, within a given organ, between the fraction of a sex steroid originating from the systemic circulation and that synthesized locally by the tissue itself ^11–16^.

Advanced imaging techniques such as positron emission tomography (PET) ^17^, mass spectrometry imaging ^15^, and fluorescently labeled steroid probes ^18^ provide valuable spatial information on steroid distribution at the tissue level. However, these methods remain constrained by limited spatial resolution and insufficient capacity to discriminate molecular sources, thereby precluding precise quantification of endogenous versus exogenous contributions. Consequently, and in the absence of more physiologically relevant alternatives, the prevailing approach continues to rely on comparing tissue steroid levels before and after administration of steroidal compounds in animals whose principal endogenous source has been either preserved or ablated by chemical or surgical castration ^19,20^. Despite its utility, this paradigm inherently disrupts systemic endocrine homeostasis and thus limits the physiological interpretability of the results.

To overcome these limitations, we developed a mass spectrometry-based strategy combining systemic administration of deuterium-labelled sex steroids with high-sensitivity GC-MS/MS analysis. This approach enables the simultaneous quantification of exogenous labelled steroids, endogenous hormones and their metabolites across multiple tissues in intact, non-castrated mice, allowing discrimination of exogenous tracers from endogenous pools.

Beyond methodological advances, this strategy provides a powerful framework to investigate steroid pharmacokinetics and tissue-specific metabolism under physiological conditions. Such information is critically important for understanding the mechanisms of action of pharmacological agents targeting steroid pathways, including anti-androgens, aromatase inhibitors and inhibitors of steroidogenic enzymes. By enabling quantitative mapping of steroid dynamics without disrupting endocrine homeostasis, this platform offers new opportunities to study drug effects on steroid metabolism and distribution in vivo, with direct relevance for pharmacological research in hormone-dependent diseases.

## EXPERIMENTAL SECTION

### Deuterium-Labeled and unlabeled Steroids

The following deuterated steroids were used: dehydroepiandrosterone-2,2,3,4,4-d_5_ (DHEA-d_5_, CAS 97453-25-3), dehydroepiandrosterone-19,19,19-d_3_ (DHEA-d_3_, CAS 721923-32-6), dehydroepiandrosterone-16,16-d_2_ (DHEA-d_2_, CAS 67034-83-7), testosterone-2,2,4,6,6,16,16,17-d_8_ (Testo-d_8_, CAS 2416308-73-9), testosterone-16,16,17-d_3_ (Testo-d_3_, CAS 77546-39-5), dihydrotestosterone-16,16,17-d_3_ (DHT-d_3_, CAS 79037-34-6), estrone-2,4,16,16-d_4_ (E1-d_4_, CAS 53866-34-5), 17β-estradiol-2,4,16,16-d_4_ (E2-d_4_, CAS 66789-03-5), 17β-estradiol-16,16,17-d_3_ (E2-d_3_, CAS 79037-37-9), 5α-androstane-3β,17β-diol-16,16,17-d_3_ (3β-diol-d_3_, CAS 79037-32-4), progesterone-2,2,4,6,6,17,21,21,21-d_9_ (Prog-d_9_, CAS 15775-74-3). They were purchased from TRC (Toronto, Canada), or CDN isotopes (Pointe-Claire Quebec, Canada). Deuterated steroids analyzed by GC-MS, but not available in standard solution in the laboratory: estrone-16,16-d_2_(E1-d_2_, CAS 56588-58-0), estrone-2,4-d_2_ (E1-d_2_, CAS 350820-16-5), 17β-estradiol-2,4-d_2_ (E2-d_2_, CAS 53866-33-4), testosterone-2,2,4-d_3_ (Testo-d_3_, CAS not available), dihydrotestosterone-2,2,4-d_3_ (DHT-d_3_, CAS not available), Δ4-androstendione-16,16-d_2_ (4-dione-d_2_, CAS not available), Δ4-androstendione-2,2,4-d_3_ (4-dione-d_3_, CAS not available), and Δ5-androstenediol-2,2,3,4,4-d_5_ (5-diol-d_5_, CAS not available).

The following unlabeled steroids, i.e., dehydroepiandrosterone (DHEA, CAS 53-43-0), dihydrotestosterone (DHT, CAS 521-18-6), estrone (E1, CAS 53-16-7), estradiol (E2, CAS 50-28-2), testosterone (Testo, CAS 58-22-0), and Δ4-androstendione (4-dione, CAS 63-05-8), were purchased from Steraloids Inc (Newport, USA). For further information, a supplementary list of deuterated steroids with their CAS numbers, commercially available, and derived from those used in this study, is available in the Supporting Information Text S1.

### Animals

Adult female and male mice (2-3 months of age) from the C57Bl/6 strain were purchased from Janvier Labs (Le Genest St Isle, France) or born in our animal facility. They were housed under controlled conditions (12 h light/dark cycle, temperature 22 ± 2°C) with ad libitum access to food and water. Both female and male mice received intra-peritoneal injection of either vehicle (100 µL of saline solution in 5 % ethanol), or vehicle spiked with a single deuterated steroid quantity (500 ng of E2-d_4_ in female mice, 1500 ng of Testo-d_3_, or 3000 ng of DHEA-d_5_ in male mice), or a mixture of three deuterated steroids at these quantities (in female and male mice) during the light phase between 09:00 and 12:00, and animals were randomized across treatment arms. The steroid doses were defined from preliminary data, considering injection safety and compound pharmacokinetics to ensure detectable tissue levels within 30 minutes after IP administration. Although mouse models usually require higher amounts of DHEA and testosterone than E2, our doses, arbitrary but adjustable, were selected to characterize tissue penetration and metabolic profiles rather than to reproduce physiological hormone ratios.

Mice were anesthetized by ketamine/xylazine to collect blood from the retro-orbital sinus, or by cardiac puncture at different time following injection, and then killed by cervical dislocation at the end of observation period (in female mice for E2 and E2-d_4_ kinetics study: 0, 1, 3, 5, 7, 10, 30, 60, 120 min; in female and male mice for a mixture of E2-d_4_, Testo-d_3_ and DHEA-d_5_ kinetics study: 0, 1, 3, 5, 7, 10 min; for E2-d_4_, Testo-d_3_, and DHEA-d_5_ tissue perfusion studies: 15, or 30 min). In females, oestrous cycle stage was not determined at the time of sampling, and inter-animal variability is expected to include a component attributable to cycle variation.

Depending on the sex of the animal, ovaries, testis, uterus, prostate, caput epididymis, seminal vesicle, brain, hippocampus, and hypothalamus were collected, and immediately frozen in liquid nitrogen and stored at-80°C. Blood was allowed to clot at room temperature for at least 15 min, and then centrifuged at 5000 g for 5 min to obtain serum. All experimental procedures were approved by the institutional animal care and use committee of the Université Paris Cité by the French Ministry of Agriculture (APAFiS #58093).

### Design of Internal Standard Solutions (IS) and Standard Solutions (STD)

Deuterated or ^13^C-labeled compounds are commonly employed as internal standards in GC-MS assays. They are added in known and identical amounts to each sample within a given analytical series, thereby allowing correction for variations inherent to instrumental performance and sample preparation steps, including extraction, purification, and derivatization. In this study, the analytes of interest included deuterium-labeled steroids administered *in vivo* and their potential metabolites across various tissues. Moreover, due to the temporary unavailability of certain deuterated derivatives in the laboratory, both the internal standard (IS) solution and the calibration ranges were specifically adapted. Details regarding the composition and preparation of IS and STD solutions are provided in tabular form in the Supplementary Tables S1 and S2.

### GC-MS Parameters

All results presented in this work were generated using modified versions of the primary GC-MS/MS analytical method previously developed for steroid quantification ^21^. For each injection, gas chromatography and mass spectrometry parameters were strictly maintained. Only the sections of the method specifying analytical targets, calibration range composition, and internal standard selection were adjusted to meet the specific requirements of each analytical series. Aged female rabbit serum, twice stripped using charcoal-dextran, served as the matrix for calibration and quality control (QC) standards. Matrix-matched calibrations for each tissue were not performed; therefore, inter-tissue matrix effects cannot be excluded.

### Sample Extraction and Purification

Briefly, each sample (serum, tissue homogenate, calibration standard, quality control, or blank matrix) was processed in an 8-mL borosilicate glass tube. Depending on the type of tissue, samples were first homogenized in a Lysing Matrix D or S tube (MP Biomedicals, Illkirch-Graffenstaden, France) containing 0.5 mL of ice-cold saline solution and 10 µL of internal standard (IS) solution, using a Ribolyser (Hybaid-RiboLyser FP120-HY-230) at a rate of two cycles of one minute each at maximum speed. The entire tube contents were then transferred by inversion and rinsed three times with 1 mL of 1-chlorobutane (HPLC-grade solvent) to ensure complete recovery. For all other sample types, the IS spiking solution was added directly to the glass extraction tube. After vortexing for 2 min and brief centrifugation, the upper organic phase was collected onto a preconditioned HyperSep SI 500-mg solid-phase extraction (SPE) minicolumn (Thermo Scientific). The column and adsorbed material were washed with 6 mL of ethyl acetate/hexane (1:9, v/v) (HPLC-grade solvents). The second fraction, containing the steroids of interest, was eluted with 4 mL of ethyl acetate/hexane (1:1, v/v) and evaporated to dryness at 60°C under a gentle stream of nitrogen.

### GC-MS/MS Derivatization and Analytical Conditions

DHEA, DHEA-d_5_, DHEA-d_3_, DHEA-d_2_, Testo, Testo-d_3_, Testo-d_8_, DHT, DHT-d_3_, 5-diol-d_5_, E1, E1-d_2_, E1-d_4_, E2, E2-d_2_, E2-d_3_, and E2-d_4_ were derivatized as previously described ^21^ using pentafluorobenzoyl chloride (PFBC; 103772-1G, Sigma-Aldrich, Steinheim, Germany). Final extracts were reconstituted in 20 µL of isooctane and transferred into 200-µL conical glass vials for GC injection. 4-Androstene-3,17-dione (4-dione), 4-dione-d_2_, 4-dione-d_3_, and Prog-d_9_ were derivatized as previously described ^21^ using 50 µL of a 1:1 (v/v) mixture of heptafluorobutyric anhydride (HFBA; Sigma-Aldrich) and anhydrous acetone. Final extracts were reconstituted in 20 µL of anhydrous n-hexane (HPLC-grade solvent) and transferred to 200-µL conical vials for injection into the GC system (GC-2010 Plus, Shimadzu) equipped with a VF-17MS capillary column (50 % phenylmethylpolysiloxane; Agilent Technologies). Detection was performed on a TQ8050 triple quadrupole mass spectrometer (Shimadzu). Compounds including DHEA, DHEA-d_5_, DHEA-d_3_, DHEA-d_2_, Testo, Testo-d_3_, Testo-d_8_, DHT, DHT-d_3_, 5-diol-d_5_, E1, E1-d_2_, E1-d_4_, E2, E2-d_2_, E2-d_3_, and E2-d_4_ were analyzed using a chemical ionization (NCI) source operated in Q3 single ion monitoring (SIM) mode, whereas 4-dione, 4-dione-d_2_, 4-dione-d_3_,and Prog-d_9_ were analyzed using an electron impact (EI) source operated in multiple reaction monitoring (MRM) mode. For NCI detection, methane was used as the reagent gas. GC was performed in pulsed splitless mode with a 1-min splitless time.

The oven temperature was initially set at 150°C for 0.5 min, increased to 305°C at 20°C/min and held for 3.6 min, then ramped to 335°C at 30°C/min and held for 1.7 min. The injector and transfer line temperatures were 290°C and 280°C, respectively. Helium was used as the carrier gas at a constant flow rate of 0.96 mL/min, and the CI source temperature was maintained at 220°C. For EI detection, GC was performed in splitless mode with a 1-min splitless time. The oven was initially held at 70°C for 1 min, ramped to 238°C at 25°C/min, then to 261°C at 5°C/min. The injector and transfer line temperatures were 290°C and 280°C, respectively, with helium carrier gas maintained at 0.70 mL/min. The EI source temperature was set at 230°C. Linearity of steroid quantification was verified by plotting the ratio of steroid peak area to internal standard peak area against steroid concentration for each calibration level. Analytical accuracy, target ions, corresponding deuterated internal controls, detection ranges, lower limit of quantification (LLOQ), and intra-and inter-assay coefficients of variation (CVs) for quality controls are summarized in Supplementary Table S3.

### Qualitative Analysis of DHEA-d_5_, Testo-d_3_ and E2-d_4_ Deuterated Solutions Injected into Mice

To determine the proportion of non-deuterated DHEA, Testo, and E2 contained in the respective deuterated steroid-concentrated solutions to be injected into the animal, i.e., DHEA-d_5_, Testo-d_3_ and E2-d_4_, respectively, a preliminary qualitative control using the conditions of method measurement was carried out in Full Scan MS (scanning between 50 and 950 amu). According to the results, the proportion of non-deuterated steroid for 100 % of deuterated steroid was 0.102 % for the DHEA-d_5_ solution, 0.070 % for the Testo-d_3_ solution, and 0.280 % for the E2-d_4_ solution. Detailed results are reported in Supplementary Data S1 to S3.

### *In vitro* Studies using deuterated E2 in AT29 cells

The AT29 cell line established from a granulosa tumour of the ovary was generously provided by Dr di Clemente (Sorbonne Université, INSERM, Centre de recherche Saint-Antoine) ^22^. We previously demonstrated that these cells are estrogen-responsive ^23^. Only cells at an early passage (<30) were used to ensure consistency in response. Cells were maintained at 37°C in a humidified atmosphere with 5 % CO2 in complete medium consisting of Dulbecco’s modified Eagle medium: nutrient mixture F-12 (DMEM/F12) (Gibco) containing 10 % foetal bovine serum (FBS) and 0.5 % penicillin/streptomycin. Cells were seeded in 12-well plates at a density of 1.0 x 10^5^ cells/well in 10 % FBS phenol red-free medium for 24 hours. Subsequently, they were treated with either vehicle control (0.1 % ethanol), E2 (Tocris) or deuterated E2 (10 nM in 0.1 % ethanol) for 24 hours. The experiment was performed in three independent biological replicates, resulting in a total of 9 samples per treatment condition. Following treatment, cells were lysed in RLT buffer (Qiagen RNeasy mini kit, #74106), and RNAs were extracted following the manufacturer’s protocol. RNA quality and concentration were assessed using a Nanodrop 2000 spectrophotometer (Thermo Scientific).

### RNA Extraction and Real-time PCR Analysis

RNAs from AT29 cells (500 ng) were reverse transcribed with the Superscript II reverse transcriptase (Thermo-Fisher Scientific). Real-time PCR were performed with primers for Growth regulating estrogen receptor binding 1 (*Greb1*) and the housekeeping gene Beta-2 microglobulin (*B2m*), designed using Primer3web, as previously described (27). Primers for *Greb1* were: forward 5’-GATTTGTTCACACCCTCAGAC-3’ and reverse 5’-GCATACTTGGACAGTCTCATACG-3’ primers (amplicon size 111 bp), and those for *B2m*: forward 5’-TGACCGGCCTGTATGCTATC-3’ and reverse 5’-GGCGGGTGGAACTGTGT-3’. For each primer pair, amplification efficiency was measured using serial dilution of cDNAs prepared from cDNA mix of control and treated samples, as described ^23^. Real-time PCR was performed with Lightcycler® 480 SYBR Green I Master and Lightcycler® 480 instrument (Roche Molecular Biomechemicals, La Rochelle, France), as previously described ^23^. All samples were run in triplicates. Quantification of *Greb1* expression was normalized to that of *B2m* and expressed as relative units.

### Statistical Analyses

All data were analyzed by Prism 10 (version 10.2.3. GraphPad Software). We used non-parametric tests when samples were not normally distributed as determined by Shapiro-Wilk normality tests. Depending on the experimental setting, we used one-way ANOVA (Tukey’s multiple comparisons test), Kruskal-Wallis or two-way ANOVA. The comparisons between groups were done with the nonparametric Mann & Whitney U test. Data are shown as scatter plot graphs with means ± SEM. A *P*-value < 0.05 was considered as significant and marked by letters for one-way ANOVA showing significant differences between means or * for *P* < 0.05 and ** for *P* < 0.01 in Student-t tests.

## RESULTS AND DISCUSSION

### Results

#### Pharmacokinetics of intraperitoneally administered E2, E2-d_4_, Testo-d_3_, and DHEA-d_5_ in intact female and male mice

To define an optimal temporal window for *in vivo* steroid tracing, we first characterized the serum pharmacokinetics of three deuterium-labeled sex steroids, i.e., E2-d_4_, Testo-d_3_, and DHEA-d_5_, following intraperitoneal (IP) administration in non-gonadectomized mice. Female mice received E2 or E2-d_4_ alone (n = 5; sampling from 1 to 120 min post-injection; Figure 1A), whereas male and female mice received a mixture of E2-d_4_, Testo-d_3_, and DHEA-d_5_ (n = 5; sampling from 1 to 10 min; Figure 1D).

**Figure 1.**
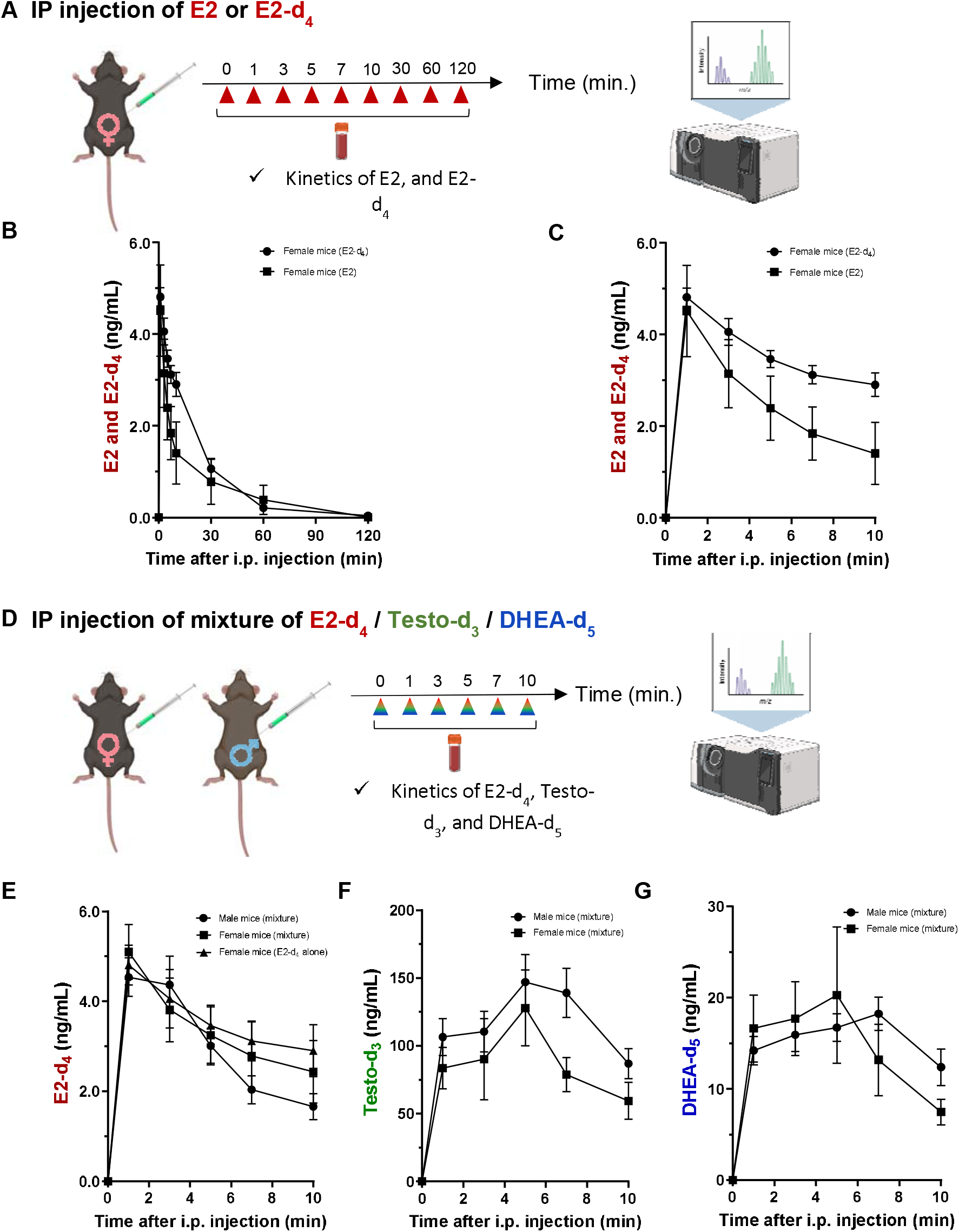
Pharmacokinetics of intraperitoneally administered E2, E2-d_4_, Testo-d_3_, and DHEA-d_5_ in non-castrated female and male mice. (A) Schematic of serum collection after E2 and E2-d_4_ injection. Created in BioRender. Wy, N. (2025) https://BioRender.com/jgbda7i. (B) Time course of E2 and E2-d_4_ following a single intraperitoneal injection at 1, 3, 5, 7, 10, 30, 60, and 120 min (n = 5 females). (C) Kinetics during the first 10 min: E2 reached a Cmax of 4.51 ng/mL (±0.45) at 1 min; estimated half-life ≈ 13 min, and E2-d_4_ reached a Cmax of 4.81 ng/mL (±0.20) at 1 min; estimated half-life ≈ 13 min. (D) Schematic of serum collection after E2-d_4_ injection. Created in BioRender. Wy, N. (2025) https://BioRender.com/jgbda7i. (E) Pharmacokinetics of E2-d_4_ following a single injection of the three-compound mixture at 1, 3, 5, 7, and 10 min (n = 5 females, 5 males). Female E2-d_4_ Cmax = 5.10 ng/mL (±0.27) at 1 min; half-life ≈ 13 min. Male E2-d4 Cmax = 4.54 ng/mL (±0.43) at 1 min; half-life ≈ 13 min. (F) Pharmacokinetics of DHEA-d_5_ after a single injection of the mixture at 1, 3, 5, 7, and 10 min (n = 5 females, 5 males). Female DHEA-d_5_ Cmax = 20.3 ng/mL (±3.3) at 5 min; half-life ≈ 12 min. Male DHEA-d_5_ Cmax = 18.2 ng/mL (±1.8) at 7 min; half-life ≈ 13 min. (G) Pharmacokinetics of Testo-d_3_ after a single injection of the mixture at 1, 3, 5, 7, and 10 min (n = 5 females, 5 males). Female Testo-d_3_ Cmax = 128.0 ng/mL (±12.5) at 5 min; half-life ≈ 11 min. Male Testo-d_3_ Cmax = 147.2 ng/mL (±20.4) at 5 min; half-life ≈ 11 min.

In female mice, E2 levels raised rapidly in blood, reaching a peak serum concentration of 4.51 ± 0.45 (SEM) ng/mL one minute after injection. For E2-d_4_ levels, the maximum serum concentration was also reached one minute after IP injection, but with a longer clearance time than that observed for E2, reaching a peak serum concentration of 4.81 ± 0.20 (SEM) ng/mL, followed by a rapid decline with an apparent half-life of approximately 13 minutes (Figures 1B and 1C).

In female and male mice, co-administration of E2-d_4_, Testo-d_3_ and DHEA-d_5_ quantities did not significantly alter the pharmacokinetic profile of E2-d_4_, which exhibited comparable peak concentrations (5.10 ± 0.27 ng/mL) and clearance kinetics in both sexes (Figure 1E). Among the three steroids, E2-d_4_ displayed the fastest systemic distribution, peaking within 1 min, whereas Testo-d_3_ and DHEA-d_5_ reached maximal serum concentrations at approximately 5 to 7 min post-injection (Figures 1F and 1G). The pharmacokinetic profile of Testo-d_3_ was similar in female and male mice, with the mean peak concentration measured 5 min after injection. However, the average Testo-d_3_ concentrations in male mice over the entire study period were 13–43 % higher than those measured in female mice under identical experimental conditions (Figure 1F). Regarding the pharmacokinetic profiles of DHEA-d5, similar patterns were observed in male and female mice. However, mean serum concentrations during the first 7 min were approximately 15% higher in females than in males, and the mean peak concentration was reached 2 min earlier in females (Cmax at 5 min in females vs. 7 min in males) (Figure 1G).

Despite lower administered doses (500 ng E2-d_4_, 1500 ng Testo-d_3_, and 3000 ng DHEA-d_5_), E2-d_4_ achieved disproportionately high serum concentrations relative to dose (5.1 ± 0.3 ng/mL at 1 min post-injection for female mice, 4.5 ± 0.4 ng/mL at 1 min post-injection for male mice). Testo-d_3_ reached the highest absolute serum concentration (128.0 ± 12.5 ng/mL at 5 min post-injection for female mice, 147.2 ± 20.4 ng/mL at 5 min post-injection for male mice), followed by DHEA-d_5_ (20.3 ± 3.3 ng/mL at 5 min post-injection for female mice, 18.2 ± 1.8 ng/mL at 7 min post-injection for male mice) (Figures 1E–1G). Inter-individual variability was observed for all three compounds; therefore, subsequent tissue concentrations were normalized to the corresponding serum levels of each labeled steroid to enable accurate inter-tissue comparisons.

#### Kinetics of E2-d_4_ incorporation in estrogen-responsive tissues of intact female mice

Prior to *in vivo* experiments, we verified that isotopic labelling did not alter estrogenic activity. In AT29 ovarian tumour cells, E2-d_4_ induced expression of *Greb1*, a canonical estrogen receptor (ER) target gene, to a similar extent as native E2 (Figure S1), indicating that deuteration does not impair ER-mediated transcriptional signalling.

Following IP administration of E2-d_4_ in intact female mice (n = 5), we quantified E2-d_4_, its primary metabolite E1-d_4_, and their endogenous counterparts in serum and classical estrogen-responsive tissues, including ovary, uterus, and hypothalamus (Figure 2A). GC-MS/MS analysis enabled unambiguous discrimination between endogenous and labeled steroids based on their distinct mass spectra (Figure 2B).

**Figure 2.**
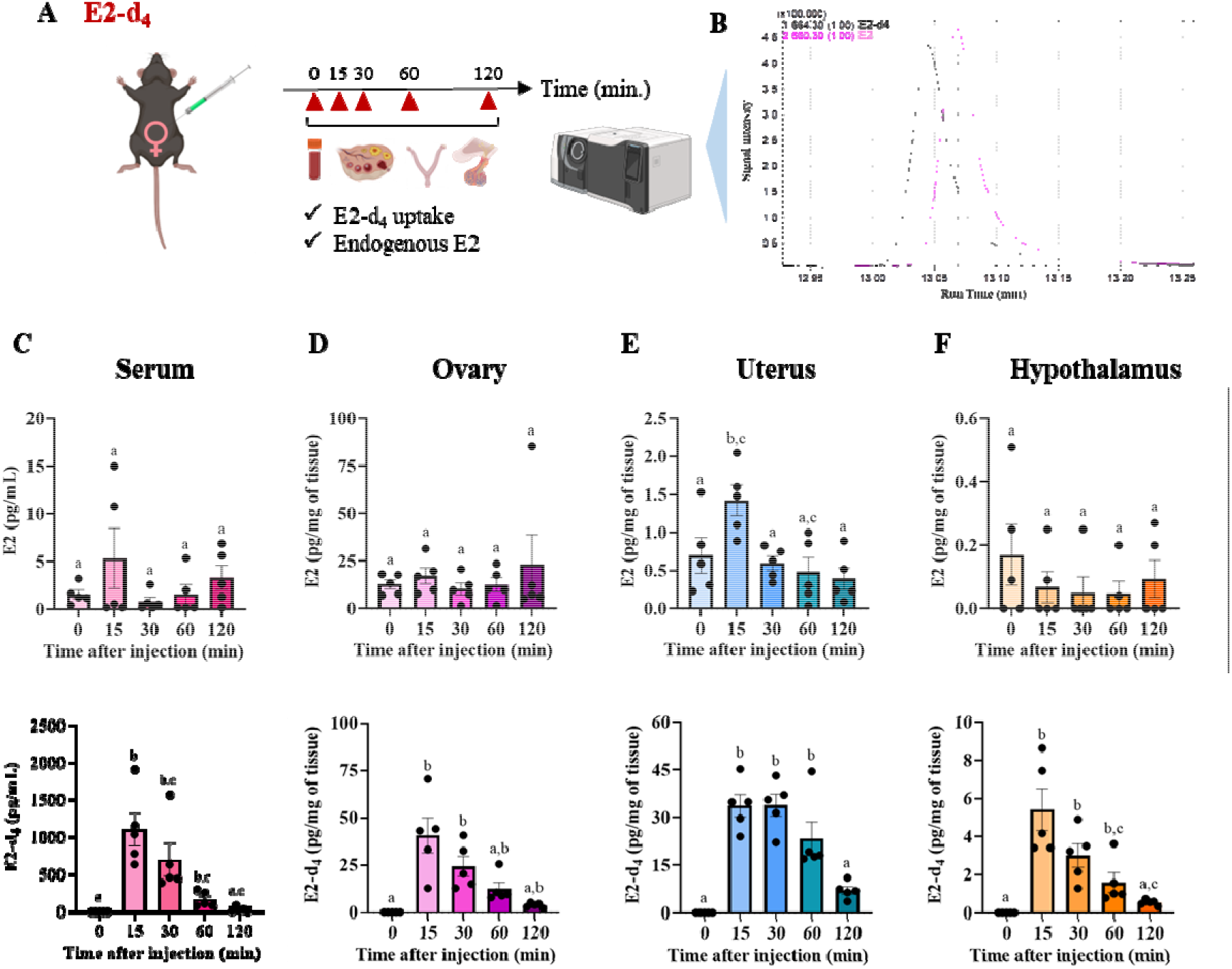
GC-MS quantification of endogenous E2 and exogenous E2-d_4_ after intraperitoneal injection in 5 adult female mice. (A) Schematic of serum and tissue collection after E2-d_4_ injection. Created in BioRender. Wy, N. (2025) https://BioRender.com/jgbda7i. (B) Example chromatogram of E2-d_4_ (50 pg, retention time 13.059 min) and E2 (56 pg, retention time 13.081 min). Identification and quantification of the target steroids were performed in single-ion monitoring (SIM) mode to allow highly sensitive, targeted detection. (C-F) Detection of E2-d_4_ by GC-MS in serum (C), ovary (D), uterus (E), and hypothalamus (F). Samples were collected at baseline (0 min, no injection) and at different time points after injection of 500 ng E2-d_4_ per female. Concentrations were calculated by dividing the measured amount of E2 or E2-d_4_ by serum volume or tissue weight. Data are shown as scatter dot plots (mean ± SEM) and analyzed by one-way ANOVA (non-parametric Kruskal–Wallis test). Different letters indicate statistically significant differences between time points (P < 0.05).

E2-d_4_ was readily detected in serum, peaking at approximately 1200 pg/mL at 15 min postinjection and declining to near-baseline levels (∼35 pg/mL) by 120 min (Figure 2C). Minor fluctuations in endogenous E2 levels were observed over time, notably in the uterus, and possibly attributed to trace non-deuterated E2 contamination (∼0.28 %) in the E2-d_4_ preparation (Figure S2).

Tissue analysis revealed rapid uptake of E2-d_4_ into ovary and uterus, reaching concentrations of ∼30–40 pg/mg within 15 min and remaining stable up to 60 min before declining (Figure 2D, 2E). Administration of E2-d_4_ induced a transient threefold increase in endogenous E2 levels in the uterus at 15 min, whereas ovarian endogenous E2 concentrations remained unchanged (Figure 2D). In the hypothalamus, a site of neurosteroidogenesis characterized by low endogenous E2 levels (∼0.1 pg/mg), E2-d_4_ was detected at lower concentrations (∼5.5 pg/mg) that closely mirrored serum kinetics (Figure 2C). These results demonstrate efficient and rapid delivery of E2-d_4_ to estrogen-responsive peripheral and central tissues in intact females.

#### Metabolism of deuterated sex steroids in intact male and female mice

To assess the ability of this analytical strategy to resolve tissue-specific steroid metabolism under physiological conditions, we next examined the biotransformation of administered deuterated steroids in intact male and female mice (n = 13). Based on known distributions of steroidogenic enzymes (Table 1), distinct metabolic profiles were anticipated (Figure 3C).

**Figure 3.**
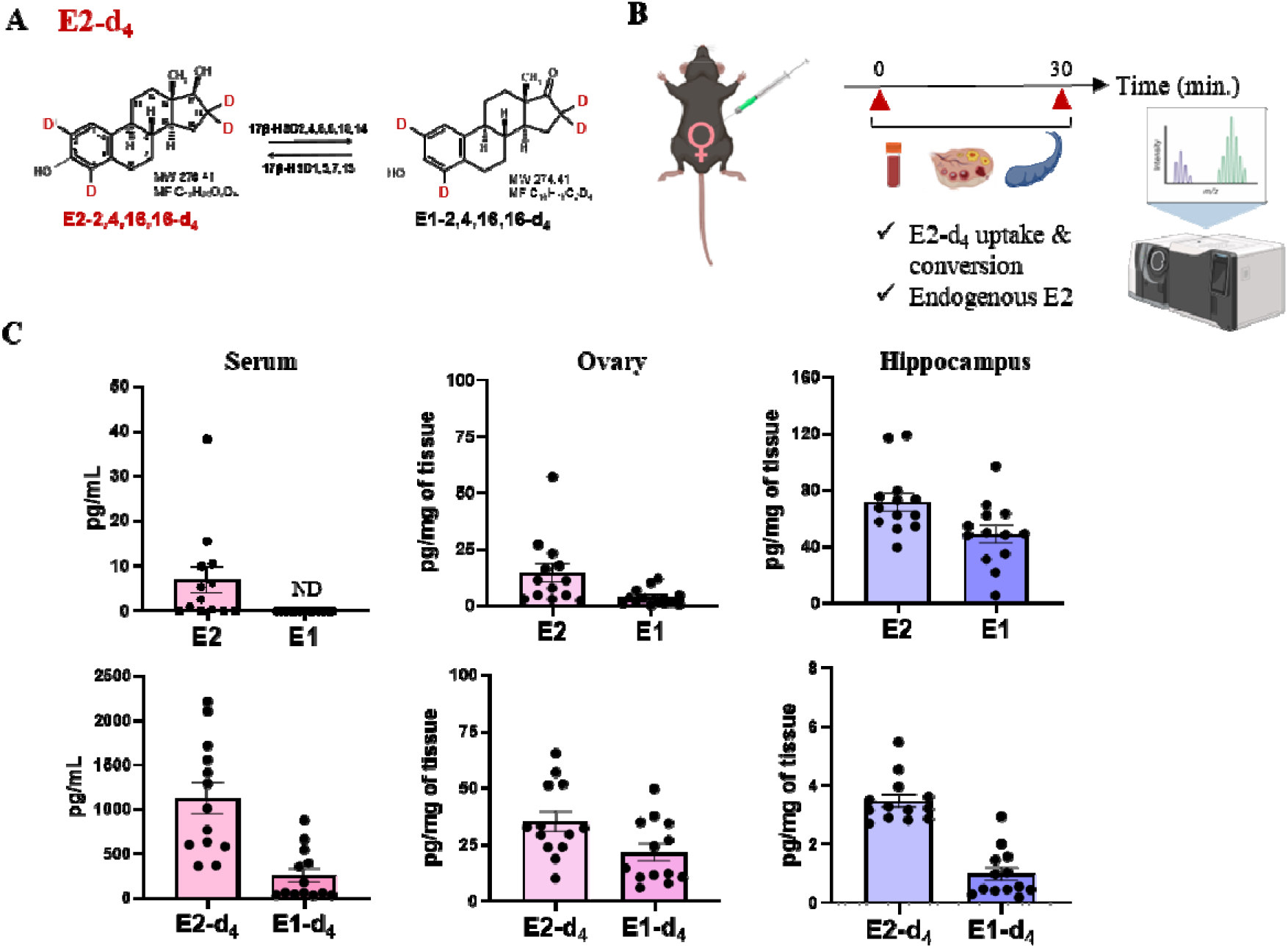
Uptake and metabolism of E2-d_4_ in the ovary and hippocampus of adult female mice. (A) Molecular weight and formula of E2-d_4_ (17β-estradiol-2,4,16,16-d_4_) and its oxidative metabolites formed by various 17βHSD isoenzymes, quantified by mass spectrometry (MW: molecular weight; MF: molecular formula). (B) Schematic of the experimental protocol. Created in BioRender. Wy, N. (2025) https://BioRender.com/jgbda7i. (C) Concentrations of E2-d_4_ and its metabolite E1-d_4_ in serum, ovaries, and hippocampus 30 min after intraperitoneal injection of E2-d_4_ (n = 13). Data are presented as scatter dot plots (mean ± SEM).

**Table 1:**
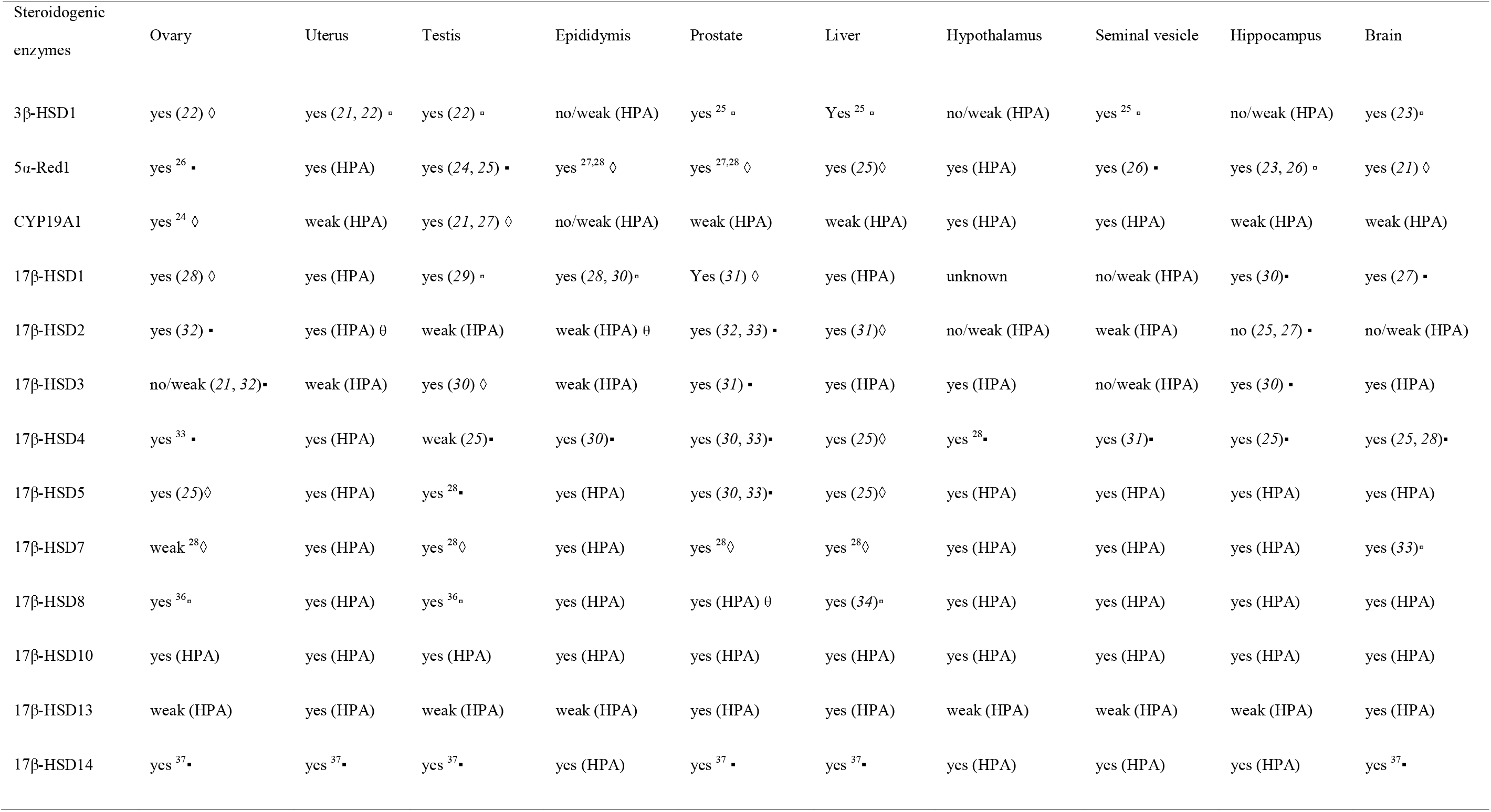
Expression of steroidogenic enzymes mRNA in human and rodent tissues (ovary, uterus, testis, epididymis, prostate, liver, hypothalamus, seminal vesicle, hippocampus, and brain). References are under bracket. HPA: Source of the website “The Human Protein Atlas”. (https://www.proteinatlas.org/): HSD3B1 https://www.proteinatlas.org/ENSG00000203857-HSD3B1/tissue SRD5A1 https://www.proteinatlas.org/ENSG00000145545-SRD5A1/tissue CYP19A1 https://www.proteinatlas.org/ENSG00000137869-CYP19A1/tissue HSD17B1 https://www.proteinatlas.org/ENSG00000108786-HSD17B1/tissue HSD17B2 https://www.proteinatlas.org/ENSG00000086696-HSD17B2/tissue HSD17B3 https://www.proteinatlas.org/ENSG00000130948-HSD17B3/tissue HSD17B4 https://www.proteinatlas.org/ENSG00000133835-HSD17B4/tissue AKR1C3 https://www.proteinatlas.org/ENSG00000196139-AKR1C3/tissue HSD17B7 https://www.proteinatlas.org/ENSG00000132196-HSD17B7/tissue HSD17B8 https://www.proteinatlas.org/ENSG00000204228-HSD17B8/tissue HSD17B10 https://www.proteinatlas.org/ENSG00000072506-HSD17B10/tissue HSD17B13 https://www.proteinatlas.org/ENSG00000170509-HSD17B13/tissue HSD17B14 https://www.proteinatlas.org/ENSG00000087076-HSD17B14/tissue

#### Conversion of E2-d_4_ to E1-d_4_ in adult female mice

Following IP injection of E2-d_4_ (500 ng), E2-d_4_, E1-d_4_, and their endogenous counterparts were quantified in serum, ovaries, and hippocampus (Figure 3C). Endogenous E2 and E1 were detected in ovaries and hippocampus, whereas serum E1 was below the limit of quantification. The ovary exhibited an approximately ninefold excess of E2 over E1, consistent with its estrogenic function, while hippocampal E1 levels were only ∼1.5-fold lower than E2.

E2-d_4_ was detected in all compartments, with substantially higher relative uptake in ovary (∼3 % of serum levels) than in hippocampus (∼0.3 %). The metabolite E1-d_4_ was also detected, with comparable E1-d_4_/E2-d_4_ ratios in ovary (∼0.5) and hippocampus (∼0.3), indicating active local interconversion of E2 to E1 in both peripheral and central tissues. These data validate the use of deuterated estradiol as a tracer for probing *in vivo* estrogen metabolism.

#### Distribution and metabolism of testosterone-d_3_ in intact male mice

Testo-d_3_ uptake and metabolism were examined 15 min after IP injection in male mice (n = 6) (Figure 4B).

**Figure 4.**
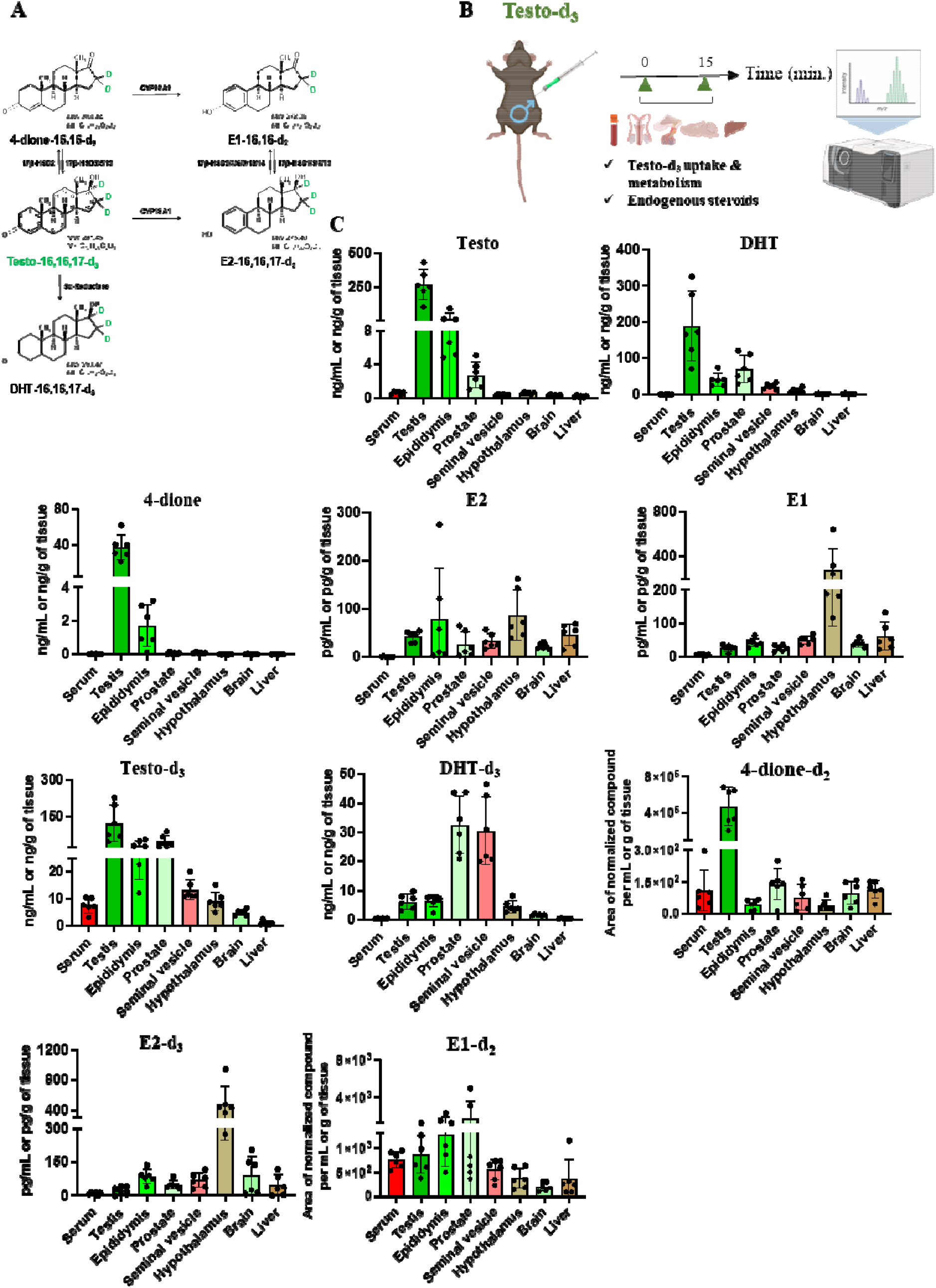
Uptake and conversion of Testo-d_3_ in reproductive and non-reproductive organs of male mice (n = 6). (A) Molecular weight and formula of major metabolites derived from Testo-d_3_ (testosterone-16,16,17-d_3_), quantified by mass spectrometry (MW: molecular weight; MF: molecular formula). (B) Schematic of the experimental protocol for Testo-d_3_ injection. Created in BioRender. Wy, N. (2025) https://BioRender.com/jgbda7i. (C) Simultaneous quantification of endogenous Testo, DHT, 4-dione, E2, and E1, and of intraperitoneally injected Testo-d_3_ and its metabolites in serum and various tissues of noncastrated male mice. Data are shown as scatter dot plots (mean ± SEM).

Endogenous and deuterated forms of testosterone, dihydrotestosterone (DHT), estrone (E1), estradiol (E2), and 4-androstene-3,17-dione (4-dione) were quantified across multiple tissues (Figure 4C).

The highest concentrations of endogenous testosterone (∼270 ng/g) and Testo-d_3_ (∼125 ng/g) were observed in testis and caput epididymis, with substantial levels also detected in prostate and seminal vesicle. Endogenous DHT was abundant in testis (∼362 ng/g) and prostate (∼69 ng/g), reflecting high 5α-reductase activity. The metabolite DHT-d_3_ was most prominent in prostate and seminal vesicle and was also detected in hypothalamus and brain, consistent with tissue-specific androgen activation.

Aromatized metabolites (E2-d_3_ and E1-d_2_) were detected in reproductive tissues and hypothalamus, with the latter exhibiting the highest E2-d_3_ levels (∼482 pg/g), supporting the importance of central aromatization of androgens in male neuroendocrine regulation.

Detection of intermediate metabolites, including 4-dione-d_2_, further confirmed rapid and tissue-dependent metabolism of Testo-d_3_ (Table 2).

**Table 2.** Putative metabolic pathways in mouse organs 15 min after IP administration of Testo-d_3_. 4-dione, Δ4-androstenedione; DHT, dihydrotestosterone; E1, estrone; E2, 17β-estradiol. Bold font indicates the preferentially produced deuterated metabolite. Epid., epididymis; Prost., prostate; sem. ves., seminal vesicle; hypot., hypothalamus. +, active; w, weakly active; +, active; w, weakly active;-, inactive. NA, not analysable due to lack of substrates.

|  | Testis | Epid. | Prost. | Sem. Ves. | Hypot. | Brain | Liver |
| --- | --- | --- | --- | --- | --- | --- | --- |
| **Deuterated metabolites** | **4-dione, E1, DHT** | DHT, 4-dione, E2, **E1** | **DHT**, 4-dione, E2, **E1** | **DHT**, 4-dione, E2, **E1** | DHT, 4-dione, **E2**, E1 | DHT, 4-dione, E2, E1 | 4-dione, E2, E1 |
| **Enzymatic activity** |  |  |  |  |  |  |  |
| CYP19A1 | w | + | + | + | + | + | + |
| 17β-HSD2 | + | + | + | + | + | + | + |
| 17β-HSD2,4, 6, 8,10,14 | + | + | + | + | w | + | + |
| 5α-Red1 | + | + | + | + | + | + | - |

#### *In vivo* conversion of exogenous DHEA-d_5_ in male mice

Finally, we investigated the distribution and metabolism of DHEA-d_5_, a major adrenal and gonadal steroid precursor, in male mice (n = 6) (Figure 5B). Fifteen minutes after IP injection, DHEA-d_5_ and six downstream metabolites, i.e., 5-diol-d_5_, 4-dione-d_3_, Testo-d_3_, DHT-d_3_, E1-d_2_, and E2-d_2_, were detected across multiple tissues (Figure 5C).

**Figure 5.**
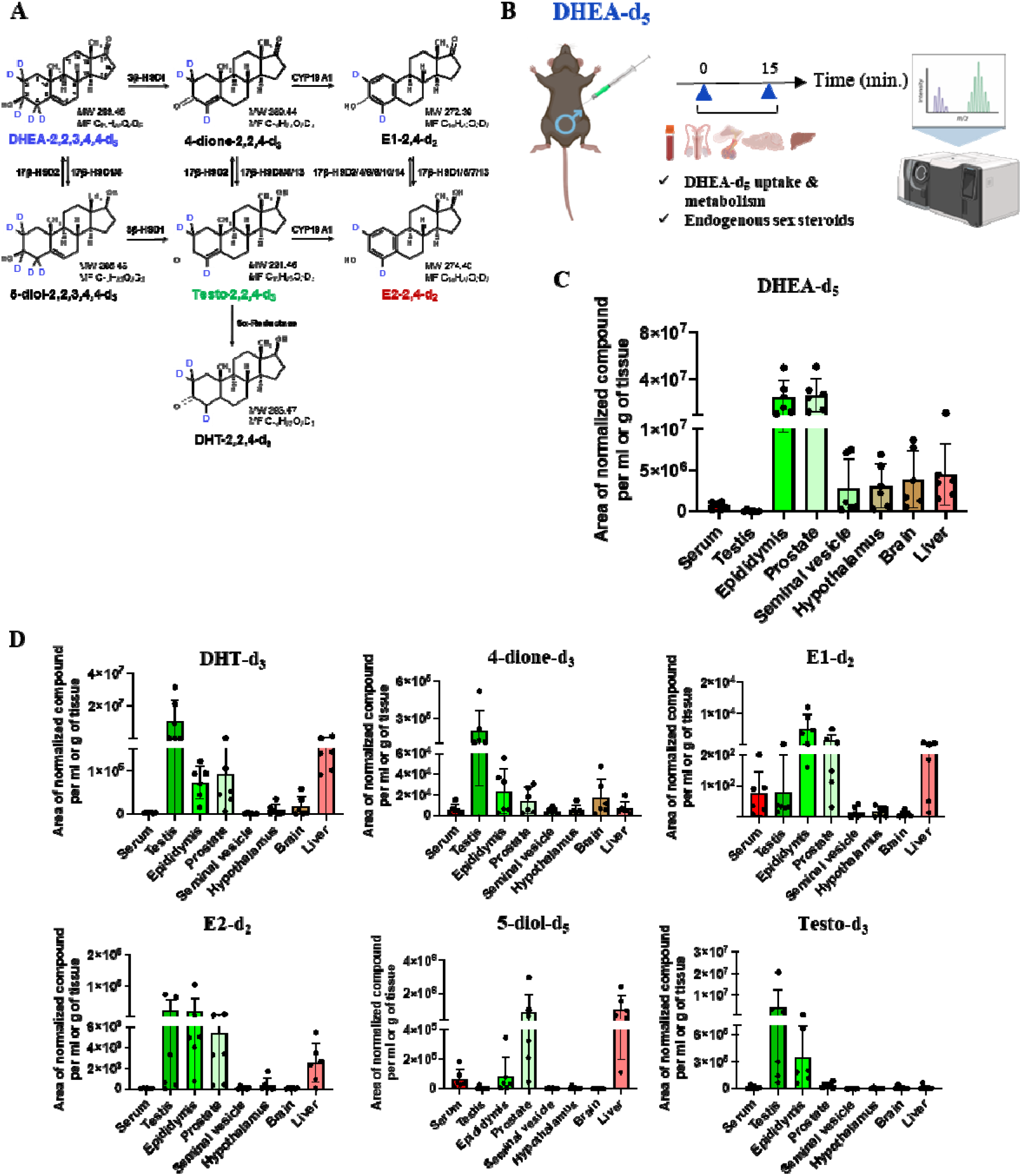
Uptake and metabolism of DHEA-d_5_ in male reproductive and non-reproductive organs (n = 6). (A) Molecular weight and formula of the major deuterated metabolites derived from DHEA-d_5_ (dehydroepiandrosterone-2,2,3,4,4-d_5_), quantified by mass spectrometry (MW: molecular weight; MF: molecular formula). (B) Schematic of the experimental protocol for DHEA-d_5_ injection. Created in BioRender. Wy, N. (2025) https://BioRender.com/jgbda7i. (C) Relative amounts of injected DHEA-d_5_ in serum and various tissues of males. (D) Relative quantities of expected deuterated metabolites detected in serum and tissues of noncastrated males 15 min after intraperitoneal injection of 3000 ng DHEA-d_5_. Data are presented as means with scatter dot plots (mean ± SEM).

DHEA-d_5_ signals were highest in caput epididymis and prostate under our assay conditions, and lowest signals observed in liver, brain, hypothalamus, and seminal vesicle, while testicular levels remained limited. Metabolite distributions reflected tissue-specific enzymatic activities: 5-diol-d_5_ was enriched in prostate and liver, 4-dione-d_3_ in testis and brain, and Testo-d_3_ primarily in testis and caput epididymis. Estrogenic metabolites were detected at lower levels across several tissues, and DHT-d_3_ was most prominent in testis and liver. These profiles delineate active and spatially organized steroidogenic pathways following DHEA-d_5_ administration (Table 3).

**Table 3:** Putative metabolic pathways in mouse organs 15 minutes after IP administration of DHEA-d_5_. 4-dione, Δ4-androstenedione; DHT, dihydrotestosterone; E1, estrone; E2, 17β-estradiol; 5-diol, Δ5-androstenediol. Bold font indicates the preferentially produced deuterated metabolite. Epid., epididymis; Prost., prostate; sem. ves., seminal vesicle; hypot., hypothalamus. +, active; w, weakly active;-, inactive. NA, not analyzable due to lack of substrates.

|  | Testis | Epid. | Prost. | Sem. Ves. | Hypot. | Brain | Liver |
| --- | --- | --- | --- | --- | --- | --- | --- |
| **Deuterated metabolites** | **4-dione, E1, E2, Testo, DHT** | **E1, E2, Testo, DHT** | **E1, E2, 4-dione, 5-diol, DHT** | / | / | 4-dione | E2, E1, DHT, 5-diol |
| **Enzymatic activity** |  |  |  |  |  |  |  |
| 3β-HSD1 | + | + | + | - | - | w | + |
| 17β-HSD1, 5 | - | + | + | - | - | - | + |
| 17β-HSD3, 5, 13 | + | + | + | NA | NA | NA | + |
| 17β-HSD1, 5, 7, 13 | + | + | + | NA | NA | NA | + |
| CYP19A1 | + | + | + | NA | NA | NA | + |
| 5α-Red1 | + | + | + | NA | NA | NA | + |

#### Impact of deuterated steroid administration on endogenous steroid levels

To determine whether administration of labeled steroids perturbs endogenous steroid homeostasis, adult male mice were injected with saline, Testo-d_3_, or DHEA-d_5_ (n = 6 per group). Endogenous sex steroids were quantified in serum and tissues 15 min post-injection (Figure 6A). Testo-d_3_ administration selectively increased endogenous testosterone levels in seminal vesicles and elevated endogenous E2 concentrations in seminal vesicles, hypothalamus, and brain, while having no significant effects in other tissues (Figures 6B to 6F). Endogenous DHT, E1, and DHEA levels remained unchanged. In contrast, DHEA-d_5_ administration did not affect endogenous testosterone, DHT, E1, or E2 levels but significantly increased endogenous DHEA concentrations in serum and liver (Figures 6B to 6F). These results indicate that acute exposure to exogenous androgens can induce rapid, tissue-specific modulation of endogenous steroid pools, consistent with localized feedback or buffering mechanisms.

**Figure 6.**
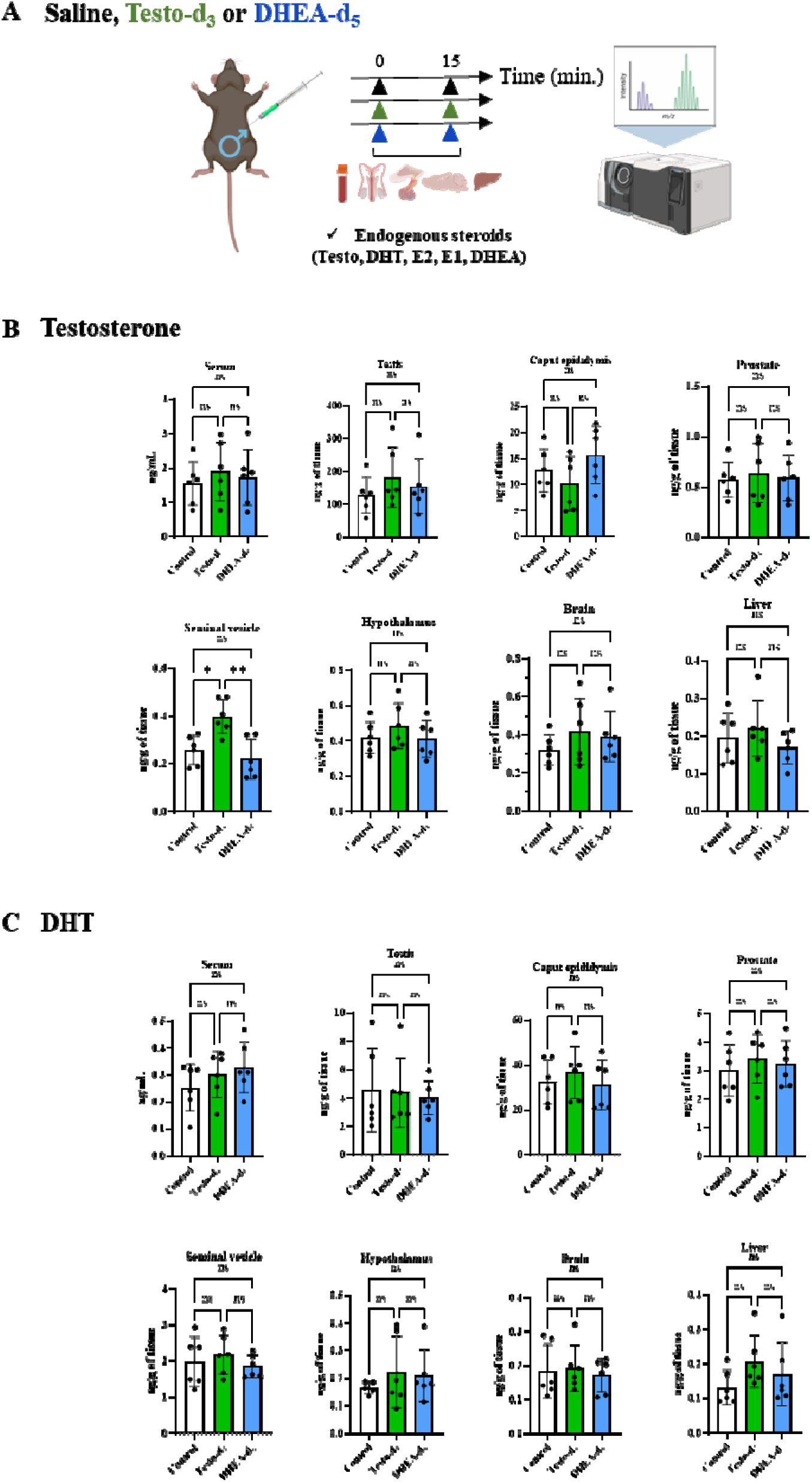
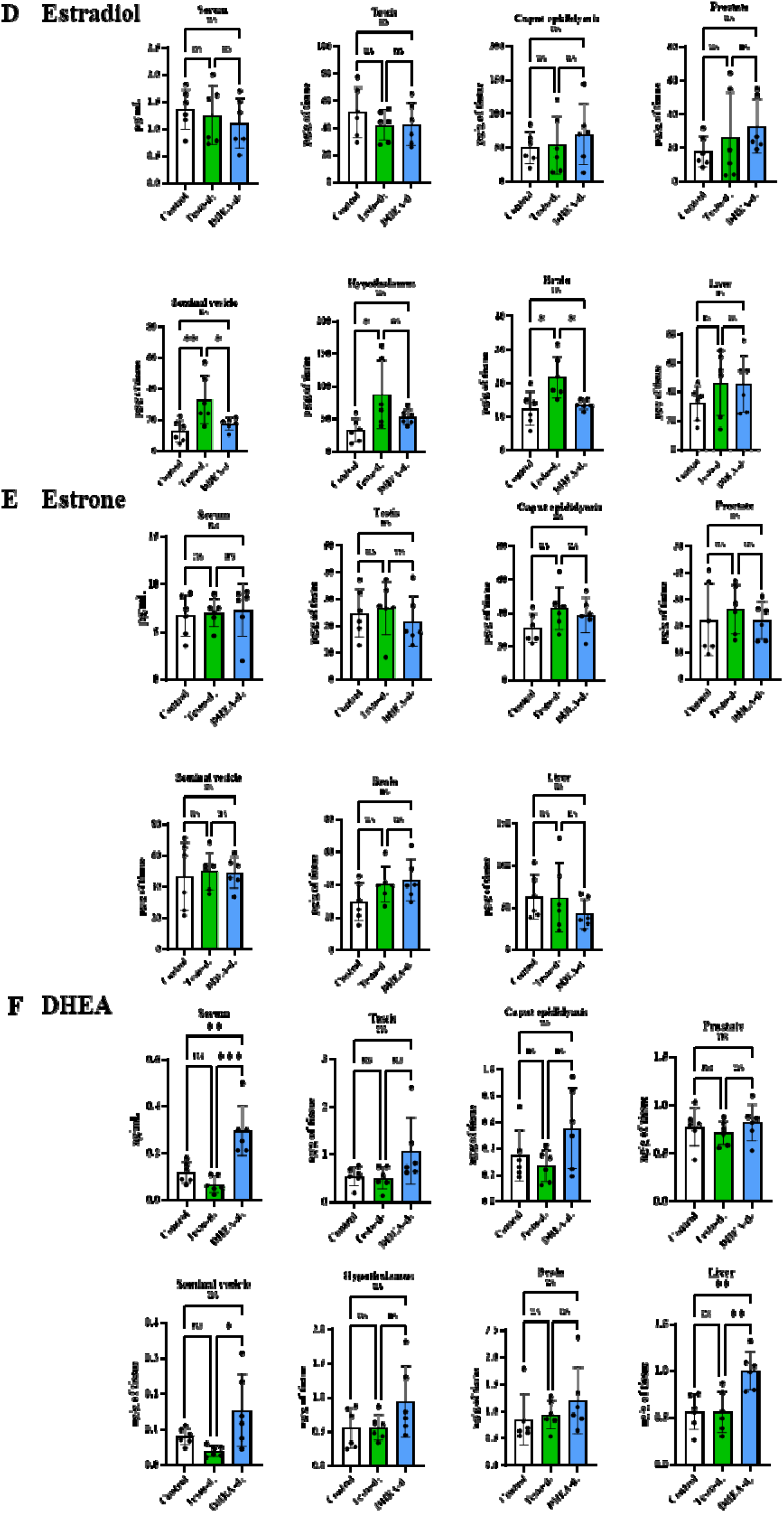
Effects of intraperitoneal injection of Testo-d_3_ and DHEA-d_5_ in non-castrated male mice (n = 6 per group). (A) Schematic of the experimental protocol for deuterated androgen administration. Created in BioRender. Wy, N. (2025) https://BioRender.com/jgbda7i. (B-F) Quantification of endogenous testo (B), DHT (C), E2 (D), E1 (E) and DHEA (F) in serum and various tissues after Testo-d_3_ or DHEA-d_5_ injection. Data are presented as means with scatter dot plots (mean ± SEM) and analyzed by one-way ANOVA; ns, not significant; *, P < 0.05, **, P < 0.01.

## Discussion

In this study, we developed and validated a mass spectrometry-based strategy combining systemic administration of deuterium-labelled sex steroids with high-sensitivity GC-MS/MS analysis. This approach enables the simultaneous quantification of exogenous tracers, endogenous hormones and metabolized steroid species across multiple tissues in physiologically intact mice. By circumventing the need for surgical castration or pharmacological suppression of steroid production, the method largely preserves systemic endocrine homeostasis over the acute time frames examined.

Our results demonstrate rapid systemic absorption and tissue-specific distribution of deuterated estradiol, testosterone and DHEA, followed by distinct metabolic conversion patterns depending on tissue type. The simultaneous detection of precursor and metabolite pools provides insights into active steroidogenic and metabolic pathways operating under native physiological conditions. These observations may reveal regulatory mechanisms that differ substantially from those observed in experimental models relying on gonadectomy or hormone replacement.

Notably, we identified marked tissue-specific metabolite accumulations, with possible conversion of E2-d_4_ to E1-d_4_ in ovary and hippocampus, testosterone-d_3_ to DHT-d_3_ in prostate and seminal vesicle, and DHEA-d_5_ to downstream androgens and estrogens in reproductive tissues and, to a lesser extent, neural tissues. These tissue distributions coincide with the expression patterns of specific enzymes summarized in Table 1, providing indirect support for a local contribution to these metabolite pools in these tissues. These findings highlight the spatial specificity of steroid metabolism and emphasize the critical contribution of local enzymatic activity to the regulation of steroid signalling. Such organ-specific metabolic patterns reinforce the concept that local steroid conversion networks play a key role in endocrine homeostasis and biological responses to circulating hormones. However, tissue accumulation of a deuterated metabolite is not, on its own, evidence of local synthesis, because labeled metabolites produced in one organ can enter the circulation and be redistributed to others. Distinguishing locally generated from systemically delivered metabolite pools will require pharmacological inhibition of specific steroidogenic enzymes, tissue-specific genetic ablation, or ex vivo tissue systems, which lie beyond the scope of the present methodological study.

Compared with traditional models based on gonadectomy or exogenous hormone replacement ^8–10^, our approach provides a major gain in physiological relevance. By maintaining endogenous hormonal feedback loops, it enables the investigation of steroid pharmacokinetics, metabolism and signalling within an intact endocrine environment. This platform therefore represents a valuable experimental framework for studying the *in vivo* regulation of steroidogenesis, tissue-specific steroid metabolism and acute steroid-mediated modulation of steroid production.

Beyond fundamental endocrine physiology, this methodology has important implications for translational and pharmacological research ^38–40^. Dysregulated steroid metabolism is a hallmark of multiple hormone-dependent diseases, including prostate and breast cancers, as well as several neurological and metabolic disorders. The ability to quantify local steroid transformations in intact animals offers the opportunity to identify tissue-specific metabolic signatures associated with disease states or therapeutic responses. In particular, this platform may facilitate mechanistic investigations of pharmacological agents targeting steroid pathways, including anti-androgen therapies ^41^, sulfatase inhibitors ^42^, aromatase inhibitors ^43^ and inhibitors of key steroidogenic enzymes ^44,45^ in animal models, including those with xenografts ^46^.

From a pharmacological perspective, the capacity to measure both circulating and locally produced steroids could significantly improve our understanding of steroid pharmacokinetics at the tissue level. Such information is highly relevant for evaluating drug efficacy, resistance mechanisms and off-target metabolic effects in therapies targeting hormonal pathways.

Furthermore, extending the approach to deuterated steroid conjugates such as DHEA-sulfate or E1-sulfate may clarify the contribution of circulating steroid reservoirs to local hormone bioavailability. Complementary LC-MS/MS analyses, particularly suited for polar conjugated steroids, could further expand the analytical coverage by enabling integrated profiling of free and conjugated steroid pools up to terminal inactivation by UDP-glucuronosyltransferases ^47^. Several technical considerations should nevertheless be acknowledged. The administered doses of deuterated steroids generated serum concentrations approximately 100-1000-fold higher than endogenous levels, reflecting a deliberate choice to ensure detection sensitivity in small tissue samples such as the hypothalamus. Consequently, the present implementation should be regarded as a pharmacological tracing approach rather than a strictly physiological one; tissue distribution and metabolite ratios may be influenced by partial saturation of binding proteins, transporters, or steroidogenic enzymes, and dose-response studies are required before quantitatively extrapolating these patterns to native steroid handling.

Temporal resolution remains limited by terminal sampling, preventing continuous monitoring of steroid kinetics within individual animals. In addition, although isotope labelling provides robust quantitative discrimination between endogenous and exogenous molecules, subtle isotope effects on enzymatic conversion cannot be entirely excluded. Future integration with spatially resolved or imaging mass spectrometry approaches may further improve temporal and anatomical resolution of steroid metabolism at cellular or subcellular scales.

In summary, this deuterium-based GC-MS/MS strategy provides a powerful and physiologically relevant approach for investigating the *in vivo* pharmacokinetics and metabolism of sex steroids. By enabling tissue-resolved mapping of steroid dynamics in intact animals, the platform establishes a foundation for mechanistic studies of endocrine regulation and offers a valuable analytical tool for research in endocrinology, neurobiology, oncology and pharmacology.

## ASSOCIATED CONTENT

All data are available in the main text or the supplementary materials.

## Supporting Information

The PDF file includes Text S1: List of deuterated steroids with their CAS numbers; Fig. S1: E2-d_4_ stimulates Greb1 expression; Fig. S2: corrected Figure 2A for circulating E2 concentrations at time 15 min; Data S1 to S3: Qualitative analysis of Testo-d_3_, E2-d_4_, and DHEA-d_5_ saline solutions; Tables S1 and S2: deuterated internal standards and ranges of standards; Table S3: GC-MS/MS Analytical Method Validation.

## Author Contributions

FG, ADV, NW, MC, CH, AI, CM, and CJG performed and analyzed the experiments. FG and CJG wrote the initial draft. FG and CJG prepared the figures. FG, ADV, and CJG edited the article. All co-authors approved the manuscript.

## Supporting information

Supporting Information_Giton et al._BIORXIV

## ACKNOWLEDGMENTS

The authors wish to thank Nathalie Di Clemente for providing AT29 cell lines, Dr Marianne Gervais-Taurel, Dr Marie Quetin, and Dr Raphaël Courcoux for their help in mouse studies, and Dr Alice Pierre for critical reading of the manuscript.

