## Supporting Information_Giton et al._BIORXIV for "Stable isotope tracing of sex steroids reveals tissue-specific steroid pharmacokinetics and metabolism in intact mice"

*Frank Giton*, Audrey Der Vartanian, Noalig Wyckens, Ioana Ferecatu, M»lanie Chester, C»line H»raud, Amine Isik, Chantal Mathis, C»line J Guigon.*

**This PDF file includes:**

Text S1

Figs. S1 to S2

Data S1 to S3

Tables S1 to S3

**Text S1: List of deuterated steroids with their CAS numbers and derived from those used in this study**

Testosterone-1,2-$d_2$, CAS 204244-83-7

Testosterone-2,2,4-$d_3$, CAS not available.

Testosterone-16,16,17-$d_3$, CAS 77546-39-5

Testosterone-1,16,16,17-$d_4$, CAS 638163-36-7

Testosterone-2,2,4,6,6-$d_5$, CAS 21002-80-2

Testosterone-2,2,4,6,6,16,16,17-$d_8$, CAS 2416308-73-9

Dihydrotestosterone-2,2,4-$d_3$, CAS not available

Dihydrotestosterone-16,16,17-$d_3$, CAS 79037-34-6

Dihydrotestosterone-2,2,4,4-$d_4$, CAS 5295-66-9

Estrone-16,16-$d_2$, CAS 56588-58-0

Estrone-2,4-$d_2$, CAS 350820-16-5

Estrone-2,4,16,16-$d_4$, CAS 53866-34-5

Estradiol-2,4-$d_2$, CAS 53866-33-4

Estradiol-16,16,17-$d_3$, CAS 79037-37-9

Estradiol-2,4,16,16-$d_4$, CAS 66789-03-5

Estradiol-2,4,16,16,17-$d_5$, CAS 221093-45-4

Dehydroepiandrosterone-16,16-$d_2$, CAS 67034-83-7

Dehydroepiandrosterone-2,2,3,4,4-$d_5$, CAS 97453-25-3

Dehydroepiandrosterone-2,2,3,4,4,6-$d_6$, CAS 1261254-39-0

4-androstene-3,17-dione-16,16-$d_2$, CAS not available

4-androstene-3,17-dione-2,2,4-$d_3$, CAS not available

4-androstene-3,17-dione-19,19,19-$d_3$, CAS 71995-66-9

4-androstene-3,17-dione-2,2,4,6,6-$d_5$, CAS 4273-44-3

4-androstene-3,17-dione-15,15,16,16,19,19,19-$d_7$, CAS not available

4-androstene-3,17-dione-2,2,4,6,6,16,16-$d_7$, CAS 67034-85-9

5-androstenediol-2,2,3,4,4-$d_5$, CAS not available

Progesterone-2,2,4,6,6,17,21,21,21-$d_9$, CAS 15775-74-3

**Figure S1**

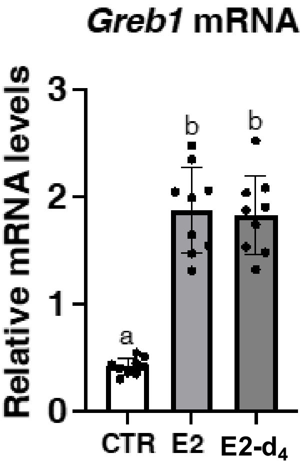


**E2-d$_4$ stimulates *Greb1* expression similarly as E2.** Relative abundance of *Greb1* transcripts in AT29 cells. Cells were treated with estradiol (E2), deuterated estradiol (E2-d$_4$) or vehicle (0.1% ethanol) for 24 hours. Relative quantification of transcripts was performed by RT-qPCR and normalization to the levels of the housekeeping gene B2m. Data are shown as scatter plot graphs with means ± SEM. Different letters indicate significant differences, with $P < 0.05$.

**Figure S2**

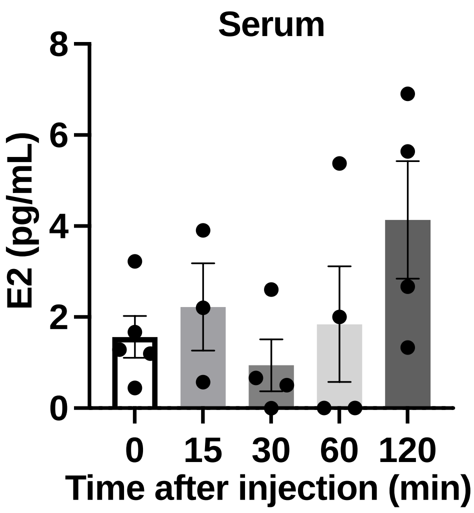


This graph shows corrected Figure 2A for circulating E2 concentrations at time 15 minutes, taking into account the presence of 0.28% unlabelled E2-d4 into the injected preparation.

**Data S1.** Qualitative analysis of Testo-d3 solution injected into mice.

| | | |
|---|---|---|
| [MS Spectrum] | TESTO-d3 | |
| # of Peaks | 794 | |
| Raw Spectrum | 11.620 (scan : 427) | |
| Background | No Background Spectrum | |
| Base Peak | m/z 485.45 (Inten : 6,124,876) | |
| Event# | 1 | |
| m/z | Absolute Intensity | Relative Intensity |
| 490.4 | 1054 | 0.02 |
| 489.35 | 5100 | 0.08 |
| 488.45 | 46092 | 0.75 |
| 487.45 | 301451 | 4.92 |
| 486.45 | 1805816 | 29.48 |
| 485.45 | 6124876 | 100 |
| 484.45 | 173243 | 2.83 |
| 483.45 | 22001 | 0.36 |
| **482.35** | **4306** | **0.07** |
| 481.35 | 1216 | 0.02 |
| 480.4 | 7374 | 0.12 |
| 479.45 | 1300 | 0.02 |
| 478.6 | 97 | 0 |
| 477.6 | 97 | 0 |
| 476.6 | 50 | 0 |
| 475.6 | 47 | 0 |
| 473.55 | 3253 | 0.05 |
| 472.45 | 6511 | 0.11 |
| 471.5 | 22377 | 0.37 |
| 470.3 | 12865 | 0.21 |
| 469.4 | 38641 | 0.63 |
| 468.25 | 4421 | 0.07 |
| 467.4 | 12719 | 0.21 |
| 466.2 | 2090 | 0.03 |
| 464.7 | 1858 | 0.03 |
| 463.35 | 1418 | 0.02 |

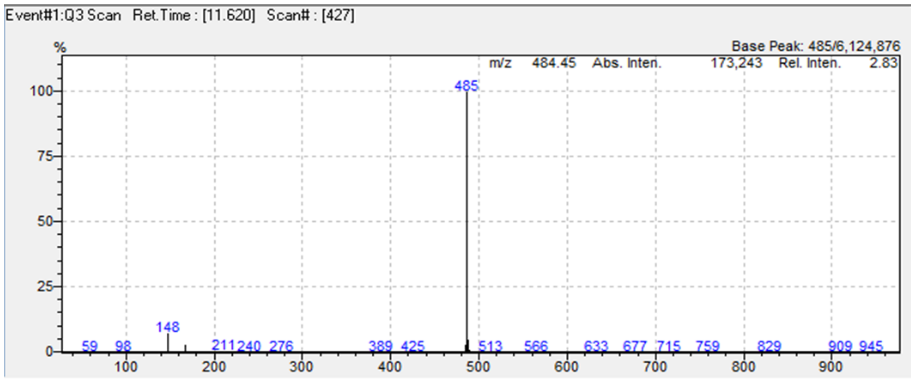


| | m/z | Ab. Intensity | % |
|---|---|---|---|
| Testo-$d_3$ | 485.45 | 6124876 | 100.00% |
| Testo | **482.35** | **4306** | **0.070%** |

**Data S2.** Qualitative analysis of E2-$d_4$ solution injected into mice.

| | | |
|---|---|---|
| [MS Spectrum] | E2-d4 | |
| # of Peaks | 783 | |
| Raw Spectrum | 13.893 (scan : 1109) | |
| Background | No Background Spectrum | |
| Base Peak | m/z 664.40 (Inten : 5,226,894) | |
| Event# | 1 | |
| m/z | Absolute Intensity | Relative Intensity |
| 670.6 | 18 | 0 |
| 669.6 | 4916 | 0.09 |
| 668.45 | 5024 | 0.1 |
| 667.5 | 74799 | 1.43 |
| 666.4 | 414689 | 7.93 |
| 665.4 | 1848849 | 35.37 |
| 664.4 | 5226894 | 100 |
| 663.4 | 188972 | 3.62 |
| 662.45 | 39080 | 0.75 |
| 661.45 | 6475 | 0.12 |
| **660.6** | **14623** | **0.28** |
| 659.5 | 8159 | 0.16 |
| 658.45 | 5954 | 0.11 |
| 655.2 | 33 | 0 |
| 653.2 | 27 | 0 |
| 652.2 | 78 | 0 |
| 651.2 | 6668 | 0.13 |
| 650.2 | 2719 | 0.05 |
| 649.35 | 23288 | 0.45 |
| 648.4 | 91588 | 1.75 |
| 647.35 | 80035 | 1.53 |
| 646.4 | 224794 | 4.3 |
| 645.35 | 177389 | 3.39 |
| 644.4 | 653868 | 12.51 |
| 643.4 | 687290 | 13.15 |
| 642.35 | 29496 | 0.56 |

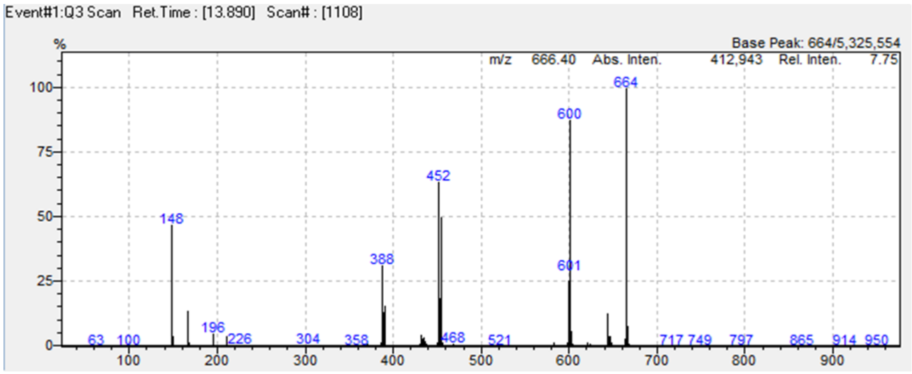


| | m/z | Ab. Intensity | % |
|---|---|---|---|
| E2-$d_4$ | 664.4 | 5226894 | 100.00% |
| E2 | **660.6** | **14623** | **0.280%** |

**Data S3.** Qualitative analysis of DHEA-$d_5$ solution injected into mice.

| | | |
|---|---|---|
| [MS Spectrum] | DHEA-d5 | |
| # of Peaks | 727 | |
| Raw Spectrum | 10.367 (scan : 51) | |
| Background | No Background Spectrum | |
| Base Peak | m/z 148.05 (Inten : 3.914.957) | |
| Event# | 1 | |
| m/z | Absolute Intensity | Relative Intensity |
| 494.5 | 455 | 0.01 |
| 492.5 | 1382 | 0.04 |
| 491.45 | 2403 | 0.06 |
| 490.45 | 23148 | 0.59 |
| 489.4 | 133137 | 3.4 |
| 488.45 | 529904 | 13.54 |
| 487.45 | 1356225 | 34.64 |
| 486.45 | 1424690 | 36.39 |
| 485.45 | 574824 | 14.68 |
| 484.45 | 40306 | 1.03 |
| 483.45 | 1874 | 0.05 |
| **482.45** | **1384** | **0.04** |
| 481.4 | 610 | 0.02 |
| 480.4 | 52 | 0 |
| 479.4 | 249 | 0.01 |
| 478.4 | 322 | 0.01 |
| 476.4 | 578 | 0.01 |
| 475.4 | 52 | 0 |
| 474.45 | 2157 | 0.06 |
| 473.55 | 2955 | 0.08 |
| 472.45 | 6966 | 0.18 |
| 471.4 | 19486 | 0.5 |
| 470.4 | 23466 | 0.6 |
| 469.45 | 19791 | 0.51 |
| 468.45 | 12683 | 0.32 |
| 467.3 | 7812 | 0.2 |

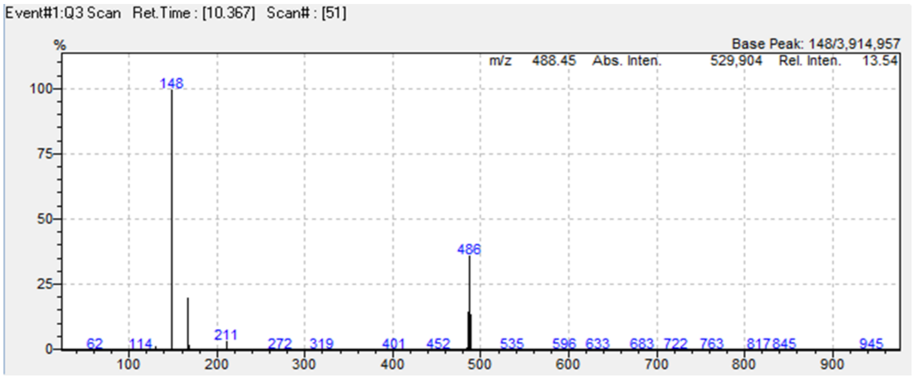


| | m/z | Ab. Intensity | % |
|---|---|---|---|
| DHEA-$d_5$ | 487.45 | 1356225 | 100.00% |
| DHEA | **482.45** | **1384** | **0.102%** |

**Table S1.** Table of deuterated internal standards and ranges of standards used to quantify the conversion of E2-d4 to E1-d4 in adult female mice.

Design of internal standard solution (IS) and standard solution (STD) for serum and tissue quantification of E1-$d_4$, E1, E2-$d_4$, and E2, after intraperitoneal injection of 500 ng E2-$d_4$

| STEROIDS | Pt0 (ng) | Pt1 (ng) | Pt2 (ng) | Pt3 (ng) | Pt4 (ng) | Pt5 (ng) | Pt6 (ng) | Corresponding IS | Quantity of deuterated steroid in 10 µl of IS solution (ng) |
|---|---|---|---|---|---|---|---|---|---|
| E1-$d_4$ | 0 | 0.001 | 0.003 | 0.009 | 0.027 | 0.081 | 0.243 | DHT-$d_3$ | 0.5 |
| E1 | 0 | 0.001 | 0.003 | 0.009 | 0.027 | 0.081 | 0.243 | | |
| **E2-$d_4$** | 0 | 0.0005 | 0.001 | 0.004 | 0.012 | 0.037 | 0.112 | 3b-diol-d3 | 1.0 |
| E2 | 0 | 0.0002 | 0.001 | 0.002 | 0.006 | 0.019 | 0.056 | | |

**Table S2.** Table of deuterated internal standards and ranges of standards used to quantify the distribution and metabolism of Testosterone-$d_3$ in intact male mice.

Design of internal standard solution (IS) and standard solution (STD) for serum and tissue quantification of DHT, DHT-$d_3$, E1, Testosterone-$d_3$, Testosterone, E2-$d_3$, E2, and 4-dione after intraperitoneal injection of 1500 ng Testo-$d_3$.

| STEROIDS | Pt0 (ng) | Pt1 (ng) | Pt2 (ng) | Pt3 (ng) | Pt4 (ng) | Pt5 (ng) | Pt6 (ng) | Corresponding IS | Quantity of deuterated steroid in 10 µl of IS solution (ng) |
|---|---|---|---|---|---|---|---|---|---|
| DHT | 0 | 0.002 | 0.006 | 0.018 | 0.054 | 0.162 | 0.486 | DHEA-$d_3$ | 2.0 |
| DHT-$d_3$ | 0 | 0.006 | 0.018 | 0.054 | 0.162 | 0.486 | 1.458 | | |
| E1 | 0 | 0.001 | 0.003 | 0.009 | 0.027 | 0.081 | 0.243 | Testo-$d_8$ | 1.0 |
| E1 -$d_2$* | 0* | 0.001* | 0.003* | 0.009* | 0.027* | 0.081* | 0.243* | | |
| **TESTO-$d_3$** | 0 | 0.03 | 0.09 | 0.27 | 0.81 | 2.43 | 7.29 | | |
| TESTO | 0 | 0.01 | 0.03 | 0.09 | 0.27 | 0.81 | 2.43 | | |
| E2-$d_3$ | 0 | 0.0009 | 0.003 | 0.008 | 0.025 | 0.075 | 0.224 | 3b-diol-$d_3$ | 1.0 |
| E2 | 0 | 0.0002 | 0.001 | 0.002 | 0.006 | 0.019 | 0.056 | | |
| 4-dione | 0 | 0.01 | 0.03 | 0.09 | 0.27 | 0.81 | 2.43 | Progesterone-$d_9$ | 4.0 |
| 4-dione-$d_2$* | 0* | 0.01* | 0.03* | 0.09* | 0.27* | 0.81* | 2.43* | | |

(*) = Proposed standard points for E1-$d_2$, and 4-dione-$d_2$. Due to the temporary unavailability of these deuterated derivatives in the laboratory, both the internal standard solution and the calibration ranges were specifically adapted.

**Table S3.** Accuracy, target ions, corresponding deuterated internal control, range of detection, low limit of quantification (LLOQ), and intra & inter assay CVs of the quality control.

| **Accuracy** (%) | **Analytes** | **Target (GC-MS) or Precursor ion analyte (GC-MS/MS)** (m/z) | **Range** (pg) | **Mean** (pg/ml) ***Intra- & Inter-assay CVs (%)*** | | | |
|---|---|---|---|---|---|---|---|
| | | | | **LLOQ** Mean Intra- & Inter assay CVs | **Low QC** Mean Intra- & Inter assay CVs | **Middle QC** Mean Intra- & Inter assay CVs | **High QC** Mean Intra- & Inter assay CVs |
| 95 - 108 | T / T-$d_3$ / T-$d_5$ / T-$d_8$ | 482.3 / 485.3 / 487.3 / 490.3 | 10 - 2 430 | 9.7 | 248.8 | 501.2 | 998.5 |
| | | | | *14.1 - 18.6* | *3.5 - 6.7* | *3.6 - 6.8* | *3.5 - 6.4* |
| 94 - 107 | E2 / E2-$d_2$ / E2-$d_3$ / E2-$d_4$ | 660.3 / 662.3 / 663.3 / 664.3 | 0.2 - 56 | 0.23 | 5.98 | 12.04 | 24.11 |
| | | | | *17.4 - 19.5* | *4.2 - 6.6* | *4.5 - 5.1* | *3.4 - 6.4* |
| 95 - 109 | DHT / DHT-$d_3$ | 484.5 / 487.5 | 2 - 486 | 1.9 | 75.2 | 148.8 | 301.2 |
| | | | | *18.1 - 19.4* | *6.5 - 8.7* | *5.1 - 7.7* | *4.6 - 6.6* |
| 93 - 108 | E1 / E1-$d_2$ / E1-$d_4$ | 464.4 / 466.4 / 468.4 | 1 - 243 | 1.2 | 25.2 | 48.9 | 100.9 |
| | | | | *18.1 - 19.7* | *6.4 - 8.9* | *5.3 - 7.9* | *4.8 - 6.8* |
| 93 - 110 | 4-dione / 4-dione-$d_2$ / 4-dione-$d_3$ | 482.20>149.20 484.20>149.20 485.20>149.20 | 10 - 2 430 | 10.3 | 252.5 | 503.4 | 1007.5 |
| | | | | *10.1 - 10.5* | *6.7 - 8.2* | *5.4 - 7.3* | *4.6 - 7.1* |
| # | Prog-$d_9$ | 519.25>519.20 519.25>147.20 | # | # | # | # | # |
| # | 3b-diol-$d_3$ | 683.4 | # | # | # | # | # |
| # | 5-diol-$d_5$ | 683.4 | # | # | # | # | # |

LLOQ : low limit of quantification; QC : quality control
